# Endophytic fungi related to the ash dieback causal agent encode signatures of pathogenicity on European ash

**DOI:** 10.1101/2023.01.04.522732

**Authors:** Maryam Rafiqi, Chatchai Kosawang, Jessica A. Peers, Lukas Jelonek, Hélène Yvanne, Mark McMullan, Lene R. Nielsen

## Abstract

Tree diseases constitute a significant threat to biodiversity worldwide. Pathogen discovery in natural habitats is of vital importance to understanding current and future threats and prioritising efforts towards developing disease management strategies. Ash dieback is a fungal disease of major conservational concern that is infecting common ash trees, Fraxinus excelsior, in Europe. The disease is caused by a non-native fungal pathogen, Hymenoscyphus fraxineus. Other dieback causing-species have not previously been identified in the genus Hymenoscyphus. Here, we discover the pathogenicity potential of two newly identified related species of Asian origin, H. koreanus and H. occultus, and one Europe-native related species, H. albidus. We sequence the genomes of all three Hymenoscyphus species and compare them to that of H. fraxineus. Phylogenetic analysis of core eukaryotic genes identified H. albidus and H. koreanus as sister species, whilst H. occultus diverged prior these and H. fraxineus. All four Hymenoscyphus genomes are of comparable sizes (55-62 Mbp) and GC contents (42–44%) and encode for polymorphic secretomes. Surprisingly, 1,133 predicted secreted proteins are shared between the ash dieback pathogen H. fraxineus and the three related Hymenoscyphus endophytes. Amongst shared secreted proteins are cell death-inducing effector candidates, such as necrosis, and ethylene-inducing peptide 1-like proteins, NLPs, that are upregulated during in planta growth of all Hymenoscyphus species. Indeed, pathogenicity tests showed that all four related Hymenoscyphus species develop pathogenic growth on European ash stems, with native H. albidus being the least virulent. Our results identify the threat Hymenoscypohus species pose to the survival of European ash trees, and highlight the importance of promoting pathogen surveillance in environmental landscapes. Identifying new pathogens and including them in the screening for durable immunity of common ash trees is key to the long-term survival of ash.

## Main

Fungal diseases have huge economic and environmental impacts worldwide. Fungal diseases infecting crops are of high research priority because they directly affect food security and the global economy. In contrast, fungal diseases of non-crop plants in natural habitats and urban areas are far less studied and, consequently, less well understood. Several plant diseases affecting environmental landscapes have caused huge permanent damage to biodiversity whilst their pathogenesis was still unknown (Fisher *et al*., 2012; Rafiqi, 2018). One such disease is ash dieback.

The ash dieback pandemic was first observed in 2002 in Poland but has quickly spread to much of Europe invading ash trees (*Fraxinus excelsior*) in natural and urban habitats (Drenkhan *et al*., 2017). The ash dieback pandemic is expected to have huge economic and environmental consequences. For example, in the UK alone, the disease is expected to cost around £15 billion (Hill *et al*., 2019). The substantial loss of *F. excelsior* is likely to negatively affect carbon sequestration and threaten around 1000 other species that are associated with ash, including birds, mammals, invertebrates, vascular plants, lichens, and fungi (Mitchell, 2014). It is anticipated that ash dieback will change the European landscape forever.

The ash dieback disease, also called Chalara dieback of ash, is caused by the ascomycete pathogen *Hymenoscyphus fraxineus* (formally known under asexual, *Chalara fraxinea*, and sexual, *Hymenoscyphus pseudoalbidus*), a commensal endophyte of Asian origin that causes no or minor symptoms when colonising leaves of Asian ash host species (Zheng and Zhuang, 2013; Zhao *et al*., 2013a; Drenkhan *et al*., 2017), yet is highly pathogenic on European ash. The genome of *H. fraxineus* is about 62 Mbp and encodes a large and diversified set of host-interacting genes (McMullan *et al*., 2018). *H. fraxineus* European population was founded by two divergent haploid individuals and has a bottlenecked genetic diversity when compared to the native Asian population of the endophyte (McMullan *et al*., 2018; Gross, Hosoya and Queloz, 2014). Subsequent introduction of new individuals is expected to increase the pathogen’s adaptive potential in Europe, which in turn would further jeopardise the survival of European ash trees. What triggers *H. fraxineus* to switch from a commensal endophyte on Asian ash to a virulent pathogen on European ash is as yet unknown.

It is widely accepted that all plants, including lichens, algae, and nonvascular plants, harbour microorganisms, called endophytes, living within them (Bacon and White, 2000). Typically, endophytes live with no apparent symptoms on plants. However, depending on environmental conditions and interactions with the host, some endophytes have visible effects on their host’s immunity and physiology. Therefore, endophytes can be pathogenic or nonpathogenic (Hardoim *et al*., 2015; Brader *et al*., 2017; Bacon and Hinton, 1996). Research in this area is scarce, although analysis of the endospheric microbiome of many plant species is increasingly detecting, through amplicon sequencing, potential pathogens that did not cause noticeable symptoms in the host at the sampling time (Pereira *et al*., 2019; Manzotti *et al*., 2020). Whether the pathogenicity of these endophytes could be triggered and how is currently unresolved. Specific ecological drivers, competition for nutrients, as well as host factors and interactions with other endophytes could all underlie the suppression of disease symptoms despite pathogen occurrence. Pathogenic fungi infecting plants in agricultural or environmental landscapes are not clustered in a particular taxonomic category that separates them from harmless endophytes. Instead, some endophytic species possess pathogenic and nonpathogenic trophic states. For example, *Sydowia polyspora* is both a foliar commensal endophyte and a preemergent seed pathogen in *Pinus ponderosa* (Ridout and Newcombe, 2017). Many fungal species include both pathogenic and non-pathogenic members. For example, *Fusarium oxysporum*, a ubiquitous soil-borne fungus, is classified in more than 150 forms or *formae speciales* of pathogenic and non-pathogenic strains, each one with the ability to infect one or a group of plant species (Nirmaladevi *et al*., 2016). Pathogenic *F. oxysporum* species often carry dispensable chromosomes that are not required for non-pathogenic growth but are virulence-associated and carry genes coding for specific host-interacting proteins and toxins, also referred to as effectors. Horizontal transfer of these chromosomes from pathogenic to a non-pathogenic recipient lineage of *F. oxysporum* renders the latter pathogenic on the respective host (van Dam *et al*., 2017). Such dispensable chromosomes have, however, not yet been identified in *H. fraxineus* (McMullan et al., 2018).

Forecasting fungal pathogen invasions in environmental landscapes is challenging and requires cross-disciplinary experimental approaches because of a myriad of variables and the complexity of interactions between the invading species and biological and physical characteristics of the recipient ecosystem (Santini *et al*., 2013; Alpert, Bone and Holzapfel, 2000; Hayes and Barry, 2008). The main factors include pathogen virulence and residence time, host susceptibility and abundance, as well as climatic factors (Hayes and Barry, 2008).

*Hymenoscyphus* is a large genus of the family of Helotiaceae and includes over 150 species (Kirk *et al*., 2008), with new species increasingly being discovered (Zheng and Zhuang, 2013; Çetinkaya and Uzun, 2020; Gross and Han, 2015). *Hymenoscyphus* species were long classified as saprotrophic decomposers and non-pathogenic endophytes up until the emergence of *H. fraxineus* as a successful invasive pathogen of ash trees in Europe. Here, we interrogate if other species of the genus *Hymenoscyphus* could be pathogenic and cause disease on European ash. We address this question by studying three *Hymenoscyphus* species closely related to *H. fraxineus*: *H. albidus*, isolated from European ash growing in Wales (UK), *H. koreanus*, and *H. occultus*; two newly-discovered species of Asian origin and not (yet) introduced to Europe; they have been isolated by Gross and Han (2015) from petioles of Korean ash (*F. chinensis* subsp. *rhynchophylla*) (Gross and Han, 2015; Kosawang *et al*., 2020).

The fungal secretome refers to the set of functionally diverse families of secreted proteins, many of which are involved in a range of diverse biological processes that directly affect the outcome of plant-fungi interactions. Examples include cell wall degrading enzymes, extracellular proteinases, toxins, and effector proteins that are upregulated during *in planta* growth of the pathogen and implicated in the suppression of host immune defences (Vincent, Rafiqi and Job, 2019). Whilst identifying and analysing predicted secretory proteins involved in disease development can’t help accurately predict pathogen invasions, it increasingly contributes to our understanding of pathogenicity and host responses. Here, we sequence and compare the genomes of three *Hymenoscyphus* species related to *H. fraxineus*. Since these fungal species can vary in their secretome predominantly by gene gains and losses or by the rapid evolution of secreted proteins, we compare the full set of proteins predicted to be secreted by these four fungal species, analyse putative effectors that could be correlated with pathogenicity and, ultimately, we test the pathogenicity of these *Hymecoscyphus* species on European ash and discuss the threat they further pose to the survival of European ash trees. The results show that in addition to the primary causative agent, *H. fraxineus*, the ash dieback disease can be caused by other member species in the *Hymenoscyphus* genus, which highlight the importance of promoting pathogen surveillance in environmental landscapes, identifying current and future threats to biodiversity, locally and at a global level.

## Results and discussion

### 1 Whole genome sequencing of three endophytic fungal species related to *H. fraxineus*

Genome sequences of three *Hymenoscyphus* species, *H. albidus* (76x coverage), *H. koreanus* (114x coverage) and *H. occultus* (102x coverage), were generated and compared to the genome of related and previously published *H. fraxineus* (McMullan *et al*., 2018). We assessed the completeness of the three *Hymenoscyphus* genome assemblies, using KAT and BUSCO. The *k-mer* distribution of such assemblies is compatible with the distribution of a complete one, and the estimated genome size is close to the actual size of the genome assembly. Approximately 96% of single copy ortholog genes from the *Sodariomycetes odb9* were completely assembled in all three genomes (Table S1). Among a total of 3725 tested BUSCO genes, only 61, 73, and 78 single-copy orthologs were missing in the *H. albidus, H. koreanus*, and *H. occultus* genomes, respectively. These results suggest that draft genome sequences generated in this study were relatively complete in terms of both assembly completeness and gene content. Genome features of *H. albidus, H. koreanus*, and *H. occultus* are detailed in Table 1.

**Table 1.** Genome assembly statistics for *Hymenoscyphus albidus, H. koreanus* F52847-4 and *H. occultus* F52847-22 isolates

| Characteristic | *H. albidus* | *H. koreanus*<br>F52847-4 | *H. occultus*<br>F52847-22 |
| --- | --- | --- | --- |
| Genome size (Mb) | 55.7 | 57.4 | 61.6 |
| No. of scaffolds | 2002 | 1517 | 728 |
| N50/L50 scaffold length (Kb) | 265/53.5 | 225/76.2 | 115/154.7 |
| Coverage | 76x | 114x | 102x |
| %GC | 44.3 | 44.1 | 42.1 |
| No. of CDSs | 13657 | 15042 | 14686 |
| Average gene length | 1520 | 1576 | 1572 |

All *Hymenoscyphus* genomes assembled here are of similar size (55-62 Mbp) and GC content (42–44%) and are comparable to *H. fraxineus*. The total number of coding sequences (CDSs) varied among the three genomes: 13,657 in *H. albidus*, 15,042 in *H. koreanus* and 14,686 in *H. occultus* (table 1). Of these CDSs, 7,791 (57.0%) in *H. albidus*, 8,313 (55.2%) in *H. koreanus*, and 8,180 (55.6%) in *H. occultus* harboured recognised protein signatures in Pfam/CDD databases. When compared to other *Hymensoscyphus* species, the predicted number of CDSs in the three genome assemblies were slightly greater than that of *H. fraxineus* (11,097 CDSs) (McMullan *et al*., 2018), but were in the same range as that of *H. varicosporoides* PMI_453 (14,929 CDSs; https://mycocosm.jgi.doe.gov/Hymvar1/Hymvar1.home.html).

OrthoMCL clustering of CDSs of all four *Hymenoscyphus* species (*H. fraxineus* and the three other related species addressed here) assigned 49,045 CDSs (90% of total CDSs) to 12,426 orthogroups (OGs), 8,665 (69.7%) of which contained CDSs from all four *Hymenoscyphus* species. Transporter, synthase, and cytochrome 450 CDSs were among the largest OGs. We found a total of 71 CDSs, representing 9 OGs specific to *H. fraxineus* (2 CDSs, 2 OGs), *H. albidus* (4 CDSs, 1 OG), and *H. occultus* (63 CDSs, 6 OGs), respectively. No CDSs specific to *H. koreanus* were found. The fact that only a small fraction, less than 1%, of predicted CDSs in all four *Hymenoscyphus* species are species-specific emphasises that these closely-related species likely share many common adaptations to similar environmental conditions. Phylogenetic analysis of core eukaryotic genes (CEGs) showed *H. albidus* and *H. koreanus* as sister species, whilst *H. occultus* was a more distant relative in the clade (Fig. 1). In previous studies, *H. fraxineus* and *H. albidus* were initially thought to be sister species, mostly due to their similar morphologies and high level of synteny (Elfstrand *et al*., 2021). In the present study we gain additional resolution with the inclusion of the newly discovered related species *H. koreanus* and *H. occultus*. Our expectation is, that with an increase in attention to this genus, additional species will be discovered and further our understanding of the biology of this group. Our finding supports the phylogenetic relatedness of *Hymenoscyphus* species previously described by (Gross and Han, 2015).

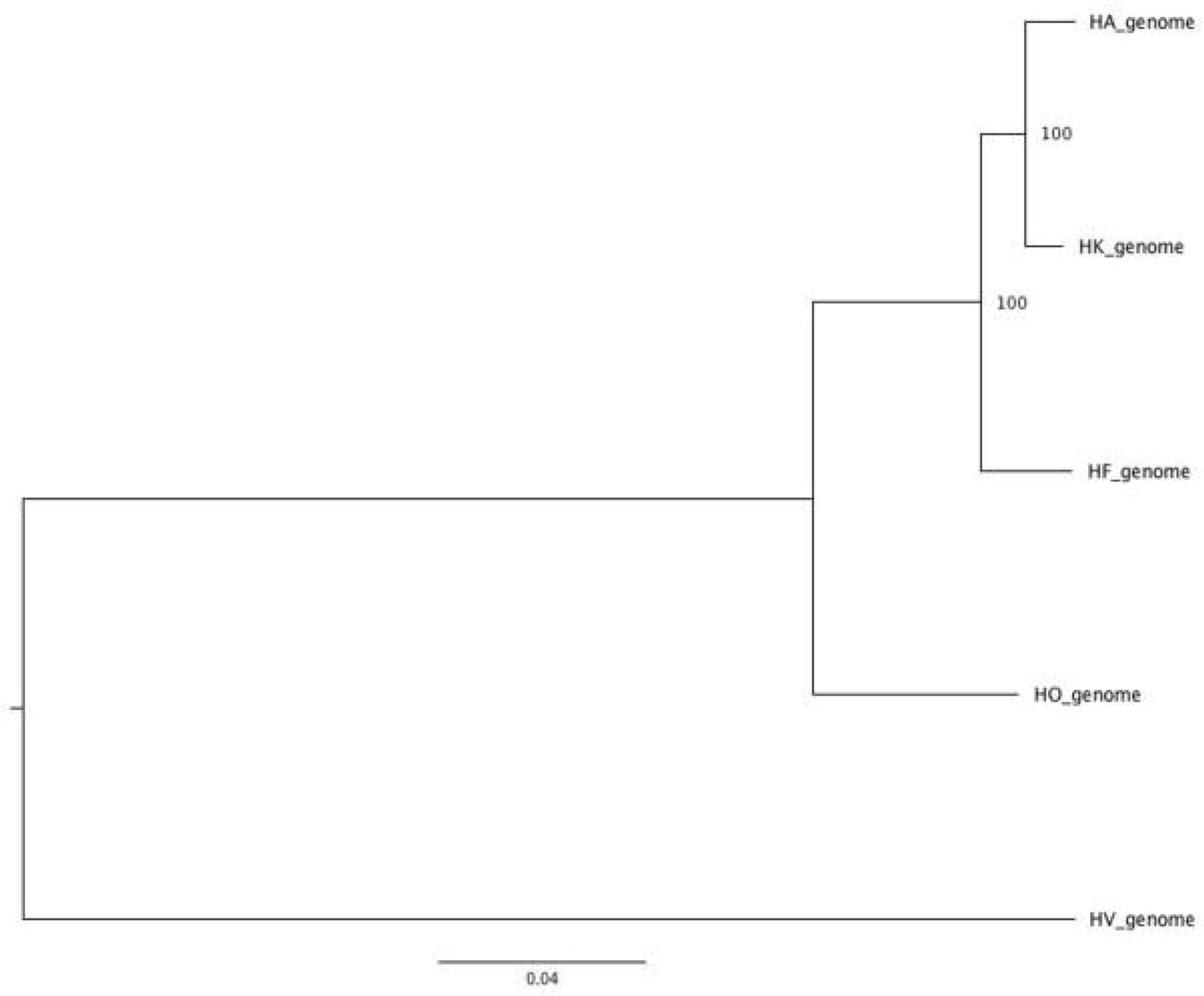

### 2 Comparative analysis of the *in silico* secretomes of *H. albidus, H. koreanus, H. occultus* and *H. fraxineus*

#### Common gene ontologies

We mined the genomes of *H. albidus, H. koreanus* and *H. occultus* for predicted secreted proteins, using a pipeline described in materials and methods. All three fungi code for secretomes of similar sizes: 1446, 1583 and 1676 predicted secreted proteins, respectively (Table S2). Secretomes of these three species were compared to that of related pathogenic *H. fraxineus*. Whilst the four predicted secretomes are highly heterogenous, strikingly a high number of proteins (1133), were shared between the ash dieback pathogen *H. fraxineus* and the three related *Hymenoscyphus* endophytes. 594 proteins were shared with *H. koreanus*, 567 proteins with *H. albidus* and 411 with *H. occultus*, with an overlap of 324 proteins shared amongst all four secretomes (Fig 2). Genes in common between pathogenic species that are involved in plant interaction shed light on a foundation of pathogenicity whereas novel genes may play a role in niche adaptation or disease development. Regardless of their phylogenetic distance, the fact that all four *Hymenoscyphus* species share a significant number of predicted secreted proteins suggests possible common infection strategies among species, which is in agreement with the observed similarity between *H. fraxineus* and *H. albidus* in developing necrotrophic growth on ash leaves (Hietala *et al*., 2022).

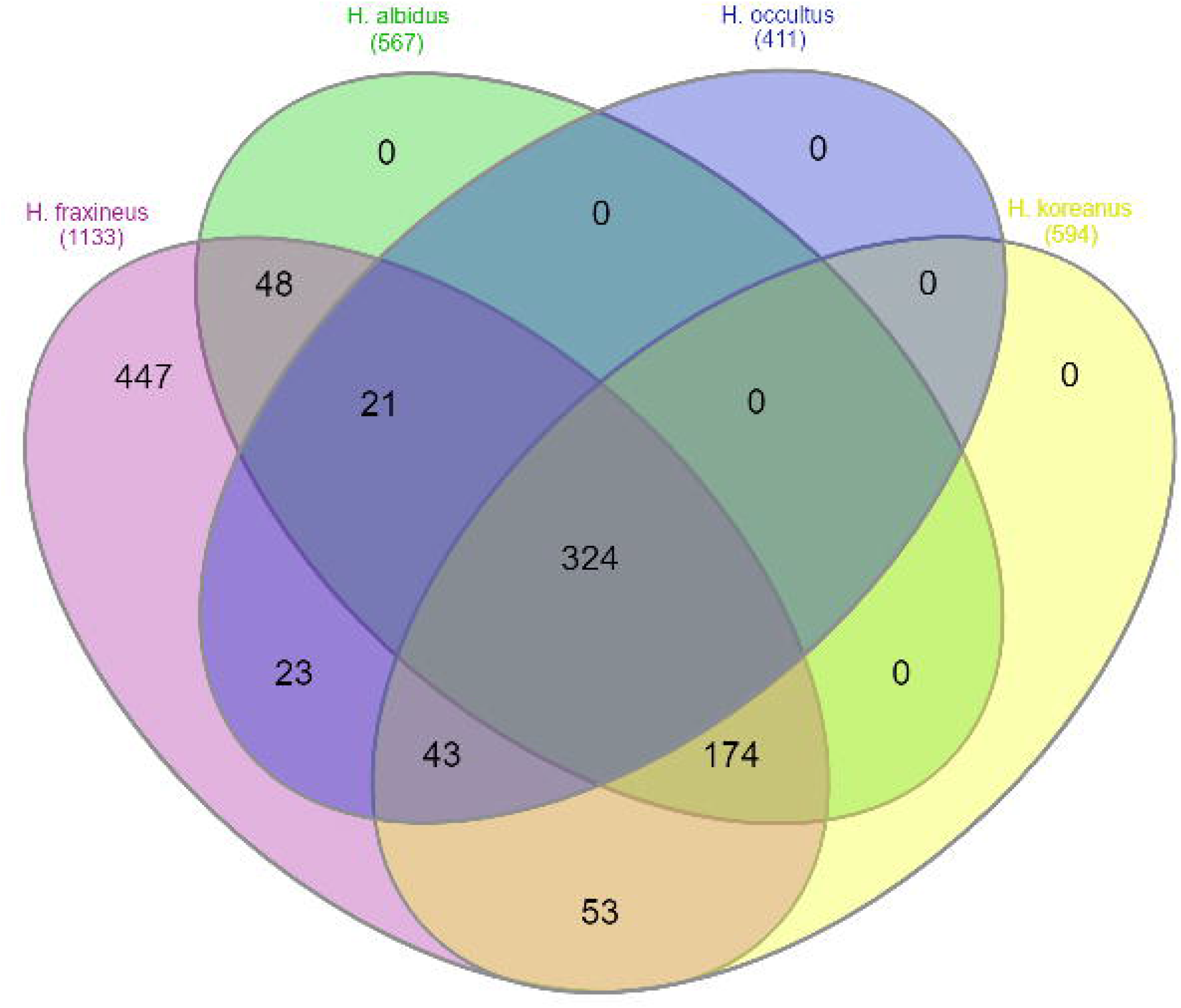

#### Polymorphism and positive selection

We analysed genetic diversity within coding sequences in all four *Hymenoscyphus* species. Genes coding for secreted proteins were associated with a SNP density that is significantly higher on average than that observed in other genes (Fig 3A). In addition, SNPs observed in secretome genes were associated with an increase in the rate of non-synonymous nucleic acid changes to the rate of synonymous nucleic acid changes (dN/dS) (Fig 3B). The polymorphism observed here in conjunction with high adaptive diversity in secretome genes among all four *Hymenoscyphus* species suggest genes coding for host-interacting proteins, including but not limited to effectors, undergo a faster rate of adaptive evolution than non-secreted genes and may play a key role in the infection process and pathogenesis on ash trees.

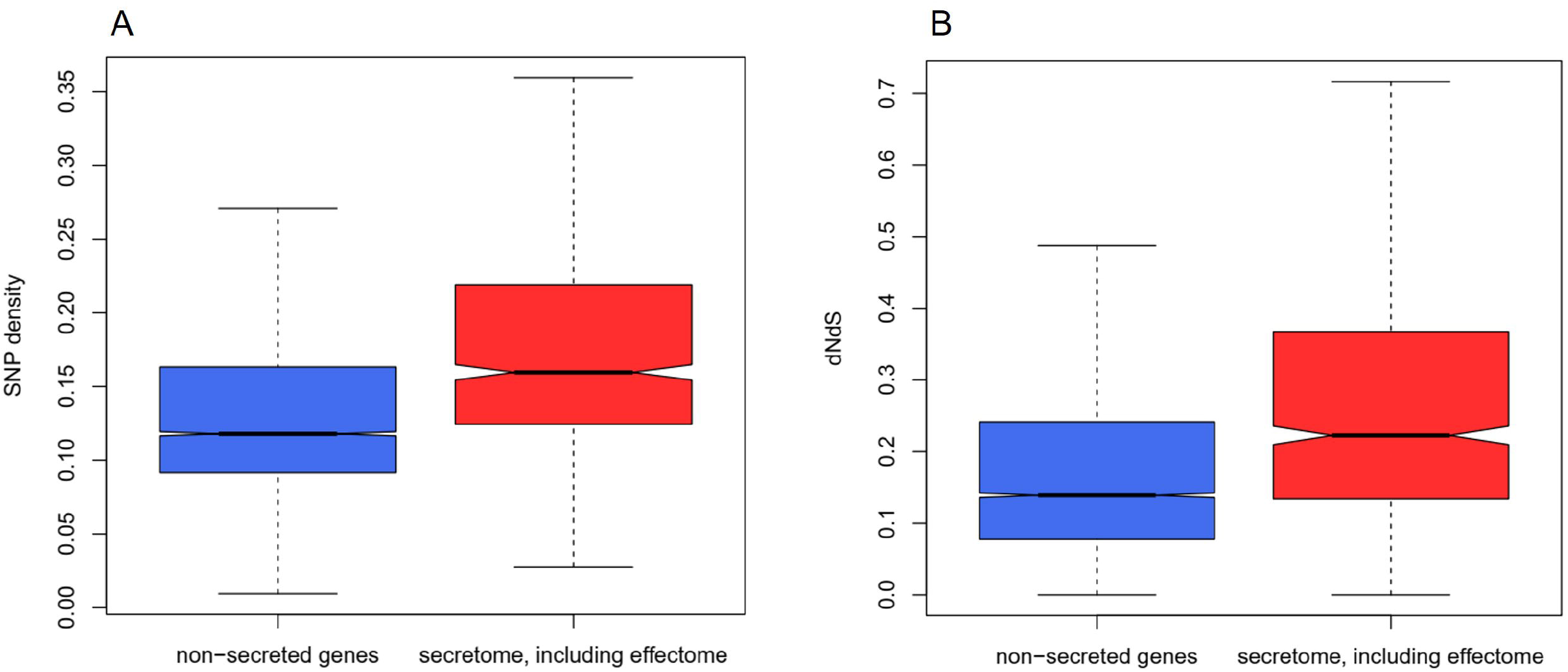

#### Large repertoires of CAZymes

To identify carbohydrate-active enzyme (CAZyme) gene content, protein sequences from the three *Hymensocyphus* species and pathogenic *H. fraxineus* were submitted for CAZymes prediction, using at least two integrated automated dbCAN2 tools (Zhang *et al*., 2018). Secretomes of all four species encode comparably high numbers (292-307) of CAZymes (Fig 4). Glycoside hydrolases (GH) appear to be the most highly represented (around 40-50%) of all CAZymes modules, followed by auxiliary activities (AA), carbohydrate esterases (CE) and carbohydrate-binding modules (CBM), respectively. CAZymes are plant cell wall degrading enzymes (PCWDs) that play a critical role in disintegrating components of the host plant cell wall, which provides pathogens with a carbon source and access to other nutrients in the host apoplast. Plant pathogenic fungi tend to secrete more CAZymes than saprophytes and symbionts. Within plant pathogens, necrotrophic and hemibiotrophic fungi contain more CAZymes than biotrophic fungi (Zhao *et al*., 2013c). The high number of CAZymes in the predicted secretomes of *Hymenoscyphus* species could, therefore, be attributed to their trophic mode (Stenlid *et al*., 2017). Cytological details of how these fungi grow *in planta* are yet unavailable, although some evidence for the formation of biotrophic fungal structures was observed in ash leaf cells inoculated with the ash dieback pathogen, *H. fraxineus*, (Mansfield, Galambos and Saville, 2018), suggesting a hemibiotrophic lifestyle, though details of how long the initial biotrophic phase takes before the pathogen switches into a necrotophic growth phase has not yet been identified (Mansfield, Brown and Papp-Rupar, 2019). Large sets of CAZymes are more frequent in necrotrophic pathogens (Zhao *et al*., 2013b). In agreement with this, a new classification of fungi based exclusively on genome-derived analysis of CAZymes has grouped *H. fraxineus* with polymertroph pathogens with a narrow host range, corresponding to necrotrophs (Hane *et al*., 2020). Indeed a recent comparative study by (Hietala *et al*., 2022) showed that both *H. fraxineus* and *H. albidus* develop similar necrotophic growth on leaves of European ash. Detailed cytological studies are further needed to understand the nature of the life style of *Hymenoscyphus* species on living ash trees and how CAZymes are involved.

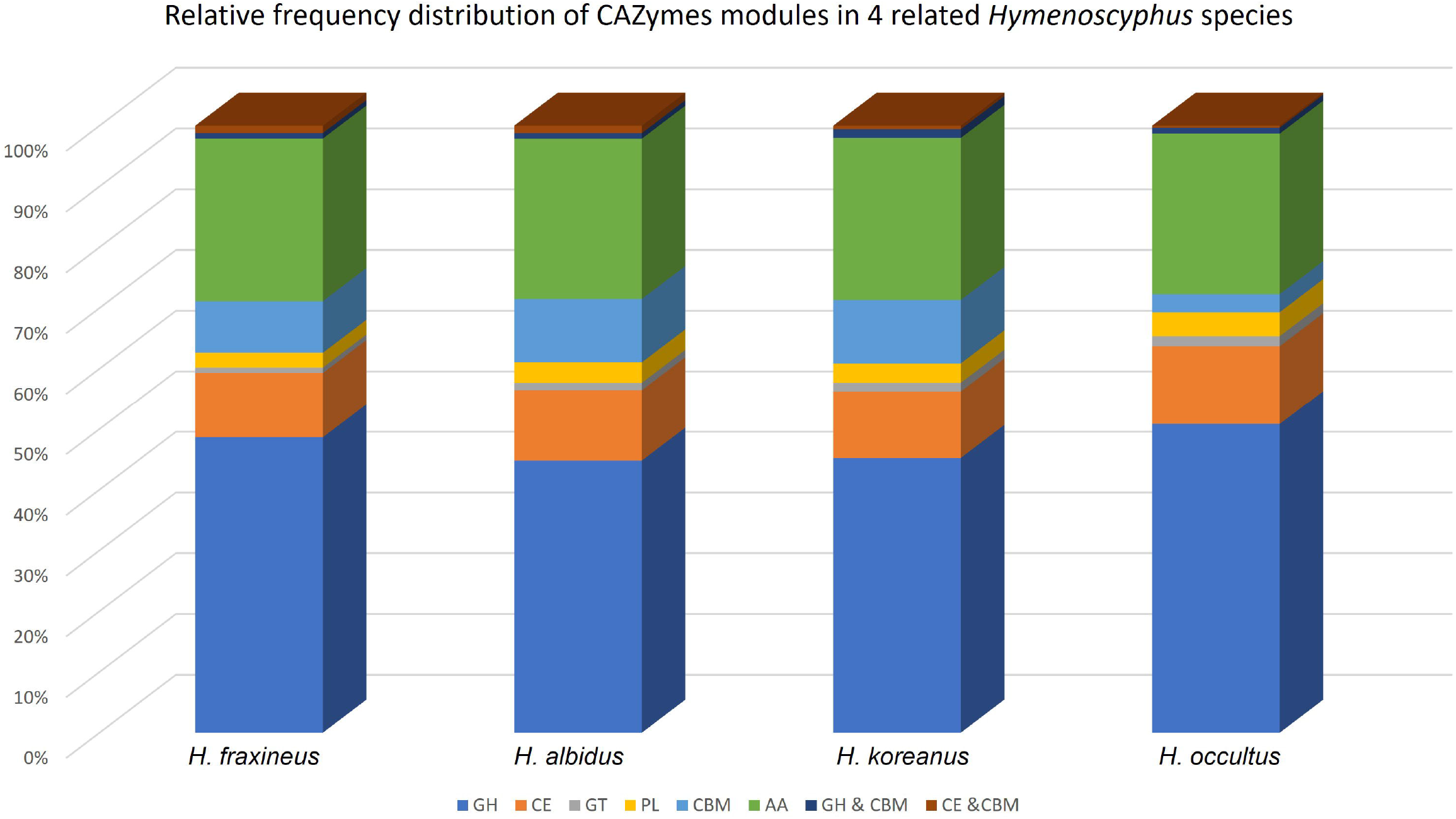

#### Shared Cell-death inducing effectors

Amongst predicted secreted proteins shared between pathogenic *H. fraxineus* and the three additional *Hymenoscyphus* species studied here are plant cell death inducing proteins. These include proteins with predicted nuclease activity. Typically, each *Hymenoscyphus* species is predicted to secrete three to four S1/P1 nucleases, one Ribonuclease 2-5A and three close homologues of T2 RNAse family protein. Secreted nuclease effectors are increasingly identified in the secretomes of fungal pathogens (Kumakura *et al*., 2021; Rafiqi *et al*., 2021; Pennington *et al*., 2019b; Rafiqi *et al*., 2022). They might either act intracellularly as a cytotoxin by scavenging host nucleic acids (Balabanova *et al*., 2012; Pennington *et al*., 2019a) or extracellularly to interact with the damage-associated molecular pattern extracellular RNA and DNA (Kumakura *et al*., 2021; Park *et al*., 2019; Widmer, 2018). Other cell-death inducing effector candidates in the secretomes of this *Hymenoscyphus* group include homologues of necrosis-inducing toxins, called Nep1-like proteins (NLPs) (Fellbrich *et al*., 2002; Bailey, 1995). NLPs cause necrosis in dicotyledonous plants and are suggested to function as virulence factors that accelerate disease development and pathogen growth in the host (Feng *et al*., 2014). Reverse transcription quantitative real-time PCR (RT-qPCR) of a selected set of genes coding for cell death inducing effector candidates showed upregulation of these genes during *in planta* growth of all *Hymenoscyphus* species (Fig 5). Predicted S1/P1 nuclease genes were induced 3dpi and 6dpi in the three *Hymenoscyphus* species as well as in pathogenic *H. fraxineus* (Fig 5A). NLP homologues are 15- and 30-fold induced in both *H. albidus* and *H. fraxineus*, respectively (Fig 5C and 5F), suggesting a potential involvement in virulence. It is particularly surprising to find that *H. albidus*, which has long been regarded as a nonpathogenic endophyte in European ash trees, is expressing PCD-inducing toxins. This is in line with recent increasing number of studies showing that *H. albidus* and *H. fraxineus*, the ash dieback primary causal agent, both develop similar necrotic growth on leaves of common ash trees. Although it causes shorter lesions on ash stems than those caused by *H. fraxineus*, *H. albidus* has been consistently shown to induce comparable lesions in leaves of European ash (*F. excelsior*), American green ash (*F. pennsylvanica*) as well as Manchurian ash (*F. mandshurica*) (Kowalski, Bilanski and Holdenrieder, 2015; Gross and Sieber, 2016; Hietala *et al*., 2022). Both expressing PCD-inducing effector candidates and causing necrosis on the host leaf tissue suggest that *H. albidus* is a mild leaf pathogen of ash trees rather than a nonpathogenic endophyte, as it has been long considered. The fact that *H. albidus* has not been associated with the recent continental breakout of the ash dieback epidemic in Europe may be attributed to its relatively low abundance, perhaps due to its low fecundity, in addition to potential environmental selection pressures that may have acted disadvantageously on its spread and virulence within its native range (Drenkhan *et al*., 2016; Hietala *et al*., 2022) as compared with invasive *H. fraxineus*. Furthermore, European ash has coexisted with native *H. albidus* for a much longer time and may, therefore, have evolved to minimise the cost of infection by this pathogen, but was in fact previously unexposed and immunologically naïve to invasive *H. fraxineus*.

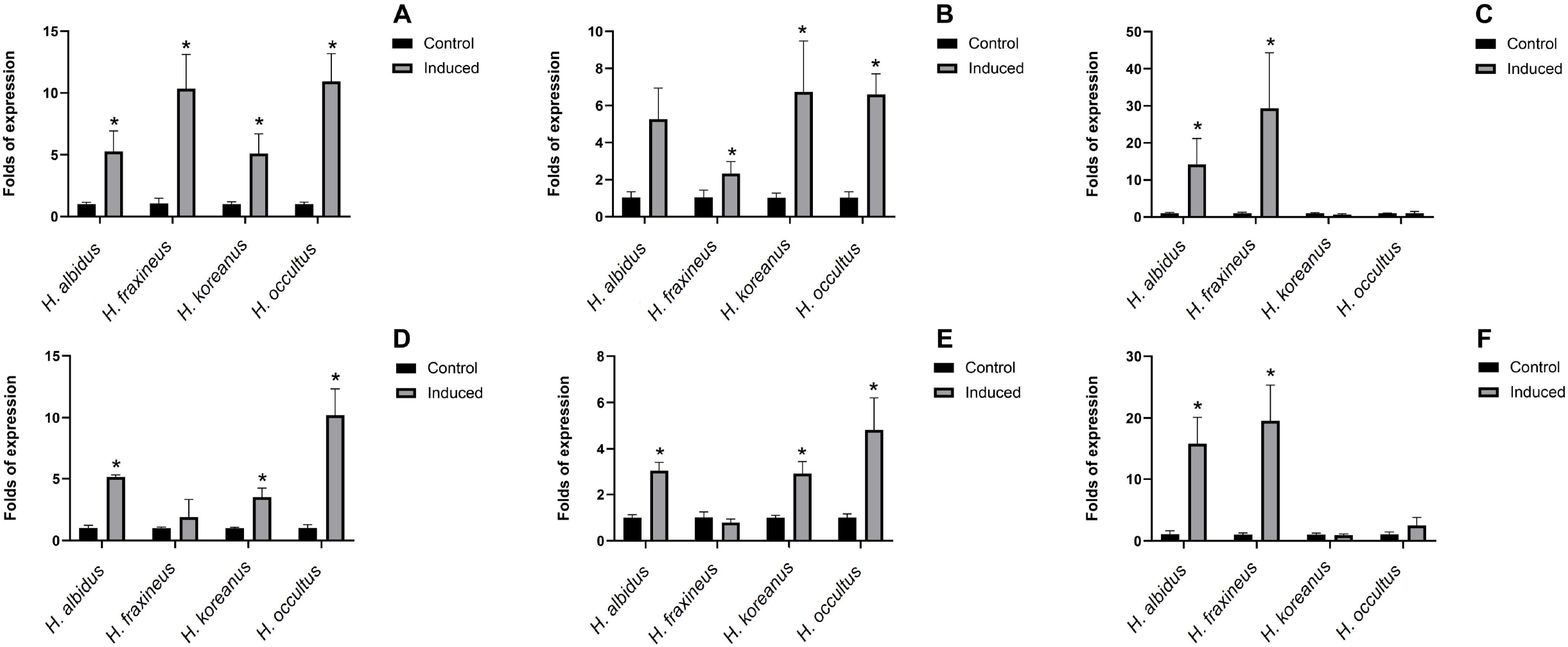

#### Pathogenicity on European ash

Even with the availability of large genome data of numerous groups of plant-interacting fungi, no study has, to date, addressed a clear correlation between genomic features or secretome composition of fungi and their pathogenicity potential. This is mainly due to the complexity of interactions between fungi and their host plants. As such, many fungal species include pathogenic as well as non-pathogenic members (Nirmaladevi *et al*., 2016), and others possess pathogenic and nonpathogenic states (Ridout and Newcombe, 2017). Therefore, predicting the pathogenicity of fungal microorganisms is as yet not accurate. Nevertheless, we set out to investigate the pathogenicity of all three *Hymenoscyphus* species, studied here. Stems of European ash seedlings were inoculated with mycelia of *H. koreanus, H. occultus* and *H. albidus*. *H. fraxineus* was used as a positive control and sterile wood plugs as negative controls. These latter did not induce the formation of any necrotic lesions on ash stems (Fig 6a-c). However, inoculations with each of the four *Hymenoscyphus* species resulted in necrotic lesions manifested by discoloration of the stem tissue to light or dark brown (Fig 6d-o). In our pathogenicity test, lesions developed by *H. fraxineus*, *H. koreanus* and *H. occultus* were relatively deeper than those formed by *H. albidus* (Fig 6m-o), suggesting a slight difference in virulence. Our results demonstrate that all four related *Hymenoscyphus* species develop pathogenic growth on European ash stems.

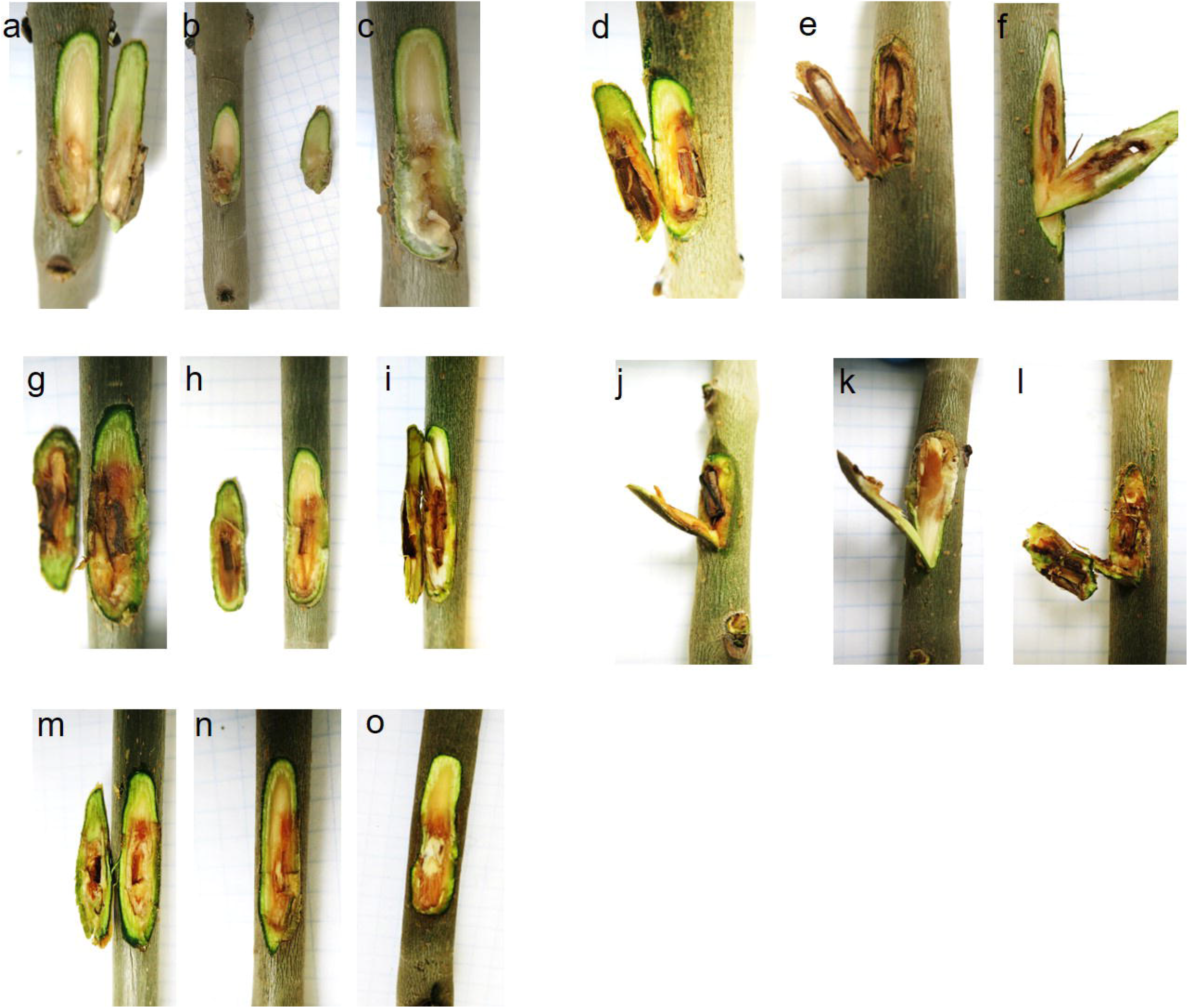

## Conclusions

In this study, we explored the pathogenicity of three fungal species closely related to the ash dieback pathogen, *H. fraxineus*. We sequenced the genomes of two new *Hymenoscyphus* species of Asian origin; *H. koreanus* and *H. occultus* as well as a native European isolate of *H. albidus*, which has long been considered as a harmless native decomposer and commensal on *F. excelsior* (Hietala and Solheim, 2011). Recently isolated from ash tree collections, the two new Asian species have not previously been reported as pathogens (Gross and Han, 2015). The genomes of all three sequenced species are of similar size and GC content and are comparable to *H. fraxineus* genome. Phylogenetic analysis of core eukaryotic genes identified *H. albidus* and *H. koreanus* as sister species, whilst *H. occultus* was a more distant relative in the clade (Fig. 1), which is in agreement with previous findings (Gross and Han, 2015). The past two decades have witnessed a surge in newly-discovered phytopathogenic species, which impacted phylogenetic analyses priorly performed. One such example is the fungal pathogen causal of sudden oak death, *Phytophthora ramorum*, which was first isolated in 2000 (Werres *et al*., 2001); fifteen years later, over 50 other Phytophthora species have been discovered and added to the Tree of Life. Currently, a large part of the inventory of pathogenic forest fungi remains undescribed; they are more likely to be discovered when they start causing noticeable, sometimes irreparable, damage to trees and the environmental landscape as a whole.

To identify genes that play a role in *Hymenoscyphus*-*Fraxinus* interaction, we analysed the secretomes of *H. albidus*, *H. koreanus*, and *H. occultus*, and we compared them to the previously described secretome of *H. fraxineus*. Whilst most predicted secreted proteins are highly heterogenous in all four secretomes, a high number of 1133 proteins were shared between the ash dieback pathogen *H. fraxineus* from one hand and the other three related *Hymenoscyphus* fungi on the other hand. One particular group of shared proteins, toxins and cell-death inducing proteins, was not only expressed, but also induced upon growth of the fungal species on detached leaves of European ash. These closely related species may, thus, share possible common niche adaptation, infection mechanisms or disease development strategies on European ash, raising the intriguing possibility that the three *Hymenoscyphus* species sequenced here may be pathogenic on European ash. Indeed, pathogenicity test using artificial infection of European ash stems with mycelia of all four *Hymenoscyphus* species produced necrotic lesions under controlled conditions, confirming the virulence of these fungal species on European ash trees (Fig 6). Lesions induced by all species were comparable, except those induced by *H. albidus*, which were slightly smaller in size and reduced in depth. Future possible introduction of Asian *H. koreanus* and *H. occultus* populations into Europe is likely to jeopardise the survival of European ash, hence the need for heightened biosecurity awareness to guide conservation strategies. This study is the first to discover new potential ash dieback pathogens, raising awareness and promoting plant disease surveillance and investigations in environmental landscapes. As a consequence of human activity, trade, and climate change, plant pathogens are broadening their geographical ranges and increasingly appearing in new habitats (Rafiqi *et al*., 2018). Forests are a particularly vulnerable ecosystem. Thus, the increasing need for pathogen surveillance and forecast for better detection and response to future outbreaks.

## Material and Methods

### Fungal isolates, maintenance, and sequencing

#### Asian isolates

*Hymenoscyphus koreanus* isolate F52847-4 and *H. occultus* isolate F52847-22 were collected in Korea from petioles of Korean ash and kindly provided by Dr. Andrin Gross (Swiss Federal Institute for Forest Snow and Landscape Research WSL, Birmensdorfm Switzerland). Fungal isolates were cultured in potato dextrose broth (VWR chemicals, USA) at room temperature and fungal mycelial mats were collected for DNA extraction 21 days after inoculation. Fungal DNA was extracted using E.Z.N.A HP plant DNA Mini kit (Omega Biotek, USA). The DNA was sent for library construction with TruSeq DNA PCR-free library prep kit with 350-bp insert (Illumina, USA) prior to sequencing with Illumina HiSeq 2500 platform on rapid run mode generating 2×250 bp (Macrogen Inc, Korea).

#### European isolates

*H. albidus* was collected near Aberystwyth in Wales, UK, and kindly provided by Dr. Fiona Corke (IBERS, Plas Gogerddan, Aberystwyth University, United Kingdom) and Dr. Anne Edwards (John Innes Centre, Norwich Research Park, United Kingdom). Single spore isolates were grown in liquid culture, harvested, freeze-dried, and the genomic DNA extracted using the MagAttract HMW DNA Kit (QIAGEN, Germany). Library preparation and sequencing were done at the Earlham Institute, Norwich. Another isolate of *H. albidus* (ID 5/37/E5) kindly provided by Dr Halvor Solheim, Norwegian Institute of Bioeconomy Research, Ås (NIBIO) was used for inoculation tests as described below.

### Genome assembly and annotation

Paired-end raw reads were adapter and quality trimmed (Q≥23) using Bbduk in the BBTools suite version 37.25 (https://jgi.doe.gov/data-and-tools/bbtools/). The draft genomes of the three *Hymenoscyphus* species were *de novo* assembled using SPAdes version 3.10.1(Bankevich *et al*., 2012) with the following parameters: k-mers 55, 77, 99 and 127, -- careful and −cov-cutoff auto. We used Blobtools version 1.1.1 (https://github.com/DRL/blobtools) and nucmer (Kurtz *et al*., 2004) to identify and dislodge scaffolds of bacterial contaminants and repetitive scaffolds (< 1,000 bp), respectively. We used the k-mer Analysis Toolkit (KAT) (Mapleson *et al*., 2017) and BUSCO version 3 (Simão et al., 2015) against the near-universal single-copy orthologues of Sodariomycetes odb9.

Annotation of the draft genomes of *H. koreanus* F52847-4 and *H. occultus* F52847-22 was achieved using the FunGAP pipeline version 1.0 (Min, Grigoriev and Choi, 2017), while the annotation of the draft genome of *H. albidus* was carried out with the funannotate eukaryotic genome annotation pipeline version 1.2 (https://github.com/nextgenusfs/funannotate). Both FunGAP and funannotate incorporate genome masking using RepeatModeler/Repeatmasker, *ab initio* gene prediction using either Augustus and Maker for FunGAP or Augustus and GeneMark for funannotate and evidence-based evaluation of the predicted gene models. Functional annotation of the predicted coding sequences (CDSs) from the *H. albidus*, *H. koreanus*, and *H. occultus* genome assemblies depended on InterproScan version 5.25.64.0 (Jones *et al*., 2014) using PFAM database version 31.0 (Mistry *et al*., 2021) for *H. koreanus* and *H. occultus* and PFAM and CDD database version 3.16 (Mistry *et al*., 2021) for *H. albidus*. The genome assemblies of *H. koreanus* F52847-1 and *H. occultus* F52847-22 and *H. albidus* were deposited at the Electronic Research Data Archive at the University of Copenhagen and can be accessed via https://sid.erda.dk/sharelink/C4VcS7pMHV.

### Phylogenetic analysis

To generate a phylogenetic tree, core eukaryotic genes (CEGs) were identified using the Ascomycota lineage in BUSCO v5.2.1 (Simão *et al*., 2015). From five *Hymenoscyphus* species, 1560 CEG single-copy orthogroups were identified. The species investigated were *H. fraxineus*, *H. albidus*, *H. koreanus*, and *H. occultus*, with *H. varicosporoides* used as an outgroup. Orthologous genes were aligned using MAFFT v7.271 (Katoh *et al*., 2002). A phylogenetic tree was generated using RaxML v8.2.12 (Stamatakis, 2014) with the PROTGAMMAGTR model and *H. varicosporoides* as the root. The tree was visualised using FigTree v1.4.3 (Rambaut, 2012).

### Secretome prediction

We used a previously described pipeline (Rafiqi *et al*., 2022) to predict fungal secretomes of *H. albidus, H. occultus*, and *H. koreanus*. The pipeline is similar to the one previously used to mine the secretome of *H. fraxineus* (McMullan *et al*., 2018), and uses SignalP 4. 1f (Nielsen, 2017) to filter proteins that contain predicted signal peptides. The set of predicted secreted proteins was further used in the pipeline to predict transmembrane helices with TMHMM 2.0c (Krogh *et al*., 2001) and cellular localization signals with TargetP 1.1b (Emanuelsson *et al*., 2000). Protein sequences containing predicted transmembrane helices or mitochondrial targeting signal were removed from the list. The remaining set of proteins was annotated with Hmmer (Zhang *et al*., 2018) against PfamScan (Finn *et al*., 2014a; Finn *et al*., 2014b) for domain information, TargetP (Almagro Armenteros *et al*., 2019) for subcellular localisation, Predictnls (Cokol, Nair and Rost, 2000) for prediction of nuclear localization signals, T-Reks (Jorda and Kajava, 2009) for detection of repeats, Disulfinder (database: uniprotkb/swiss-prot) for prediction of the disulfide bond, and MOTIF search for the search of known motifs. The positions of the motifs RxLR, [LI]xAR, [RK]CxxCx12}H, [YFW]xC, YxSL[RK], G[IFY][ALST]R, DELD, and [SG]PC[KR]P were identified with a script based on regular expressions.

### Genetic diversity

To investigate genetic diversity within coding sequences, OrthoFinder v.2.2.6 (Emms and Kelly, 2019) was used to identify orthogroups from four *Hymenoscyphus* species: *H. fraxineus*, *H. albidus*, *H. koreanus*, and *H. occultus*. Single-copy orthogroups were selected and orthologous genes from the four species were each combined into a single file. Genes were aligned using GUIDANCE2 in codons with the MAFFT algorithm (Sela *et al*., 2015). Due to erroneous sequence lengths, 37 genes were removed, leaving 7308 genes.

Synonymous (dS) and non-synonymous (dN) mutations were quantified using (Yang, 2007) PAML v4.9 with the yn00 method on the single-copy orthogroups. The dN/dS ratio was calculated. DnaSP (Rozas *et al*., 2003) was run on the orthologous genes to investigate single nucleotide varient density per gene, which was calculated by dividing the number of SNPs by the net number of sites. DnaSP failed to run on two genes, leaving 7306 genes.

### Stem inoculation

An inoculation experiment was performed on 4/7/2018 to determine if inoculation with the fungi (*H. koreanus* F52847-4, *H. occultus* F52847-22, and *H. albidus* 5/37/E5) could cause the development of necrotic lesions on *Fraxinus excelsior*. Two-year-old open-pollinated offspring from clone #35 from the clonal seed orchard Tuse Næs (N55° 45 57.99 E11° 42 47.48) that had been kept under disease-free conditions were used for the experiment. The seedlings were inoculated with infected wood plugs. Before inoculation, sterile wood plugs (3 × 4 × 10 mm^3^ in size) had been placed on five week old cultures allowing the mycelium to colonize the wood plugs. An incision was cut into the stem of the plant and the infected wood plug was placed in the incision. Parafilm was wrapped around the stem. A culture of *H. fraxineus* strain 18.3 (Kosawang *et al*., 2020) was used as a positive control and sterile wood plugs as negative controls. Three biological replicates were performed for each inoculation type (i.e. the three species and the positive and negative controls). During the experiment, the plants were kept in a closed chamber at room temperature and 16 hours of light. The experiment was terminated after 5 weeks (8/8/2018). Parafilm was removed and photos of the lesions were taken for documentation.

### qRT-PCR of selected *Hymenoschyphus* effector candidates

Six, two-week-old mycelia plugs of the three *Hymenoscyphus* species and *H. fraxineus* strain 18.3 were grown on detached leaves of common ash (*Fraxinus excelsior*) clone 27x rinsed with running tap water for 30 mins. The plugs were placed face-down allowing the fungal mycelia to interact with the leaves. Three of the plugs were harvested after 3 and 6 days after inoculation (DAI) for RNA extraction. For the control treatment, the plugs were grown in malt extract broth and were collected accordingly after 3 and 6 DAI. Total RNA was isolated from the plugs using an E.Z.N.A Plant RNA kit (Omega Bio-tek, USA) with on-column DNase I digestion, and cDNA was prepared from 500 ng total RNA using a qScript cDNA synthesis kit (Quanta Biosciences, USA) following the manufacturer’s instruction. Three biological replicates of control and treatment were performed for each species of *Hymenoscyphus*.

Gene expression analysis was performed in two technical replicates for each biological replicate on a Mx3005P qPCR system (Stratagene, USA) and FIREPol EvaGreen qPCR Mix Plus (Solis Biodyne, Estonia). Analysis of melting curves was carried out at the end of each run to detect nonspecific amplifications. Relative expression of the effector-encoding genes was calculated in relation to the expression of the beta-tubulin gene according to the 2^-ΔΔ^CT method (Livak and Schmittgen, 2001). Gene expression data were analysed statistically using student t-test with a confidential level of 95% implemented in Microsoft Excel version 16.30 (Microsoft Corp., USA).

## Data availability

The genome assemblies and annotations of *H. koreanus* F52847-1, *H. occultus* F52847-22 and *H. albidus* were deposited at the Electronic Research Data Archive at the University of Copenhagen and can be accessed via https://sid.erda.dk/sharelink/DljEqJhxMG.

## Acknowledgements

We thank Dr Andrin Gross (The Swiss Federal Institute for Forest, Snow and Landscape Research, Switzerland), Dr Fiona Corke (Aberystwyth University, Aberystwyth, UK), Dr Anne Edwards, Professor Allan Downie (John Innes Centre, Norwich, UK) and Dr Halvor Solheim (Norwegian Institute of Bioeconomy Research, Ås (NIBIO)) for providing fungal material. We thank our funders Godfred Birkedal Hartmanns Familiefond (2017), the Danish Council for Independent Research (grant no. 6111-00254) and Royal Botanic Gardens Kew Pilot Study Fund 11231-105. Work at the Earlham Institute (EI; MM, JAP & HY) was supported by Biotechnology and Biological Sciences Research Council (BBSRC), part of UK Research and Innovation, through the Core Capability Grant (BB/CCG1720/1) and (BBS/E/T/000PR9818) WP1 Signatures of Domestication and Adaptation. Sequencing at the EI is supported by BBSRC National Capability in Genomics and Single Cell Analysis (BBS/E/T/000PR9816) by members of the Genomics Pipelines and Core Bioinformatics Groups. HY & <u>JAP</u> were also supported by the BBSRC funded Norwich Research Park Biosciences Doctoral Training Partnership grants BB/M011216/1 & BB/T008717/1 as well as JAP Year in Industry (BBS/E/T/000PR9811).

## Authors contribution

MR, MM and LRN planned and designed the research. MR, CK, JAP, JL, HY, MM and LRN performed experiments and analysed data. MR, CK, MM and LRN wrote the manuscript. MR and CK contributed equally.

